# IL-1RA controls the development of esophageal cancer by inhibition of VEGF via PI3K/NF-κB signaling pathway

**DOI:** 10.1101/623637

**Authors:** Zhimin Shen, Shaobin Yu, Lei Gao, Zhun Liu, Peipei Zhang, Tianci Chai, Sui Chen, Mingqiang Kang

**Affiliations:** Department of Thoracic Surgery, Fujian Medical University Union Hospital, Fuzhou 350001, China; Key Laboratory of Ministry of Education for Gastrointestinal Cancer, Fujian Medical University, Fuzhou, 350001, China; Fujian Key Laboratory of Tumor Microbiology, Fujian Medical University, Fuzhou 350122, China

**Keywords:** IL-1RA, ESCC, VEGF, PI3K/NF-KB

## Abstract

IL-1RA has been reported to function as tumor suppressor in a variety of tumors, but its function and mechanism in esophageal cancer remains largely unknown. Our previous studies have shown that IL-1RA is downregulated in primary esophageal cancer(EC), and this downregulation of IL-1RA is closely related to TNM staging and survival prognosis. In this study, we observed that over-expression of IL-1RA inhibits proliferation, migration, and tumor growth in esophageal cancer cells(ESCC), inhibits the expression and secretion of VEGF, and inhibits tumor angiogenesis. Further studies have shown the overexpression of IL-1RA inhibits the transmission of PI3K/NF-kappaB. Taken together, this study shows that IL-1RA can regulating the development of esophageal cancer by inhibiting VEGF expression and secretion via inhibiting PI3K/NF-kappaB signaling, and it may serve as a potential prognostic marker and therapeutic target for EC.

## 1. Introduction

Esophageal cancer is the sixth most common gastrointestinal cancer in the world with the highest incidence rate and the fourth highest mortality rate[1]. In Southeast Asia, The most common type of esophageal cancer is ESCC[2]. Although current treatments, such as surgery and chemotherapy, have progressed esophageal cancer treatment, the 5-year overall survival rate of ESCC is still low (20%-30%) due to metastasis, recurrence, and resistance to chemo-radiotherapy[3]. Therefore, there is an urgent need to study the reasons behind the pool progression of esophageal cancer, as this will provide an effective strategy for the diagnosis, treatment and prognosis of esophageal cancer.

At present, studies have shown that various cytokines such as interleukin (IL), chemokines and lymphokines are involved in inflammation and its regulation, and play important roles in affecting tumor biology[4]. Inflammation participates in the construction of the tumor microenvironment by altering the homeostasis of the tumor tissue. The interaction of various cytokines secreted by tumor microenvironment infiltrating cells constitutes a complex network of cytokines, which promotes tumor progression through various factors such as pro-inflammatory, immune editing, and immune escape[5]. The related research between inflammation and cancer focuses on the promotion of cancer in the early stage[6], for example, inflammation in liver cancer can directly promote the proliferation and survival of tumor cells, and affect the surveillance of the immune system by affecting immune regulation. In addition, inflammation can also induce angiogenesis and genomic instability to promote tumor development[7]. Pro-inflammatory cytokines are key regulators of the tumor microenvironment, controlling tumor cell proliferation, promoting inflammation, angiogenesis, and tumor metastasis. In addition, the studies about IL-1β, IL-11, IL-17, IL-18, TNF, etc. have also shown that networked pro-inflammatory cytokines are key regulators of tumor microenvironment and can control tumor cell proliferation and promote Inflammation, angiogenesis, and tumor metastasis[8].

Hypoxia in the tumor microenvironment(TAM) not only induces immunosuppression but also induces angiogenesis and inflammation. Coordination and reinforcement between inflammation and angiogenic cytokine expression links, VEGF and COX2 via hypoxia-inducible and NF-κB signaling pathways[9][10]. In addition, other pro-angiogenic or pro-inflammatory factors (including TNF, IL-1, IL-6 and IL-8 /CXCL8) are expressed by HIF signaling, regulate hydrogen peroxide production and NF-κB signal. VEGF as a chemoattractant of macrophages also leads to inflammation and immunosuppression, which usually come from TAM. MMPs and inflammatory factors, especially M-CSF/CSF1 can increase VEGF expression, so it can be in inflammation / blood vessels Establish a positive feedback loop between generation and immunosuppression[11].

PI3K is an intracellular phosphatidylinositol kinase with serine/threonine (Ser/Thr) kinase activity localized on the cell membrane, and NF-κB factor is localized in the cytosol and isolated by IκB molecules. In the classical PI3K/NFκB signaling pathway, receptors located outside the cell membrane can accept various external stimuli, such as proinflammatory Cytokines, TLRs, Bacterial LPS, etc. Under stimulation conditions, IκB undergoes ubiquitination and degradation, which makes NF-κB Activation by phosphorylation, NF-κB dimer translocates into the nucleus and binds to the gene promoter of downstream proteins, exerting its functions of promoting inflammation, tumor proliferation and metastasis.[12].

Interleukin-1 (IL-1) usually play as a pro-inflammatory chemokine that can promote tumor cell proliferation and differentiation through interaction with it’s specific receptors site in the surface of tumor cell membranes.[13]. IL-1RA, IL-1α and IL-1β are all the members of the interleukin-1 family. IL-1α and IL-1β can bind to the same receptor and bear similar biological functions, such as promoting tumor growth and metastasis through enhanced angiogenesis. This is accomplished by modulating the expression of angiogenic factors, such as vascular endothelial growth factor (VEGF)[14]. IL-1RA is the biological physiological inhibitor of IL-1 which blocks the activation of IL-1 receptor via competitive binding to IL-1 with its receptor. IL-1RA is associated with a variety of diseases including type 2 diabetes, cardiovascular disease, cancer and joint disease[15]. IL-1RA inhibits angiogenesis by blocking the activation of the IL-1a / PI3K/NF-κB pathway in human colon cancer cells producing IL-1[16]. Downregulation of IL-1RA is associated with metastatic potential of gastric cancer by blocking IL-1α/VEGF signaling pathway[17]. Our previous study also showed that IL-1RA is down-regulated in esophageal cancer and is closely related to tumor TNM staging. The proliferation function of esophageal squamous carcinoma cells overexpressing IL-1RA was significantly decreased in vitro, and the expression of VEGF was decreased. However, the specific mechanism of how IL-1RA causes VEGF changes and thus affects tumor progression remains unclear. Based on the previous research results, this study further improved the related functional experiments, and further explored whether IL-1RA regulates VEGF through PI3K/NF-κB signaling pathway, and then regulates the progression of esophageal cancer.

## 2. Materials and methods

### 2.1 Cell lines and cell culture

Human EC cell lines KYSE410 and Eca109 were purchased from Hunan Fenghbio Biological Ltd, China. 293T cell was gifted from Key Laboratory of Ministry of Education for Gastrointestinal Cancer, Fujian Medical University. All the cell lines were cultured in the RPMI-DMEM (Gibco) medium supplemented with 10% fetal bovine serum (Gibco, USA). All the cell lines were maintained in an humidified incubator atmosphere with 37°C, 5% CO2.

### 2.2 Quantitative reverse transcription PCR (RT-PCR)

Extraction of total RNA from cultured cells or frozen tissues using Trizol reagent (Ambion, Carisbad, CA, USA), and strand complementary DNA synthesis using miScript Reverse Transcription Kit (Qiagen, Hilden, Germany). Real-Time PCR was performed using SYBR Premix EX Taq kit (Takara, Shiga, Japan). The specific primers were used to detect the relative mRNA expression of IL-1RA by the 2 - ct method. GAPDH is used to normalize to measure the level of expression. All the primers were designed by BioSune Biotechnology Co., Ltd (Shanghai).

### 2.3 Western blotting

Tissues or cells were lysed on ice in Western and IP cell lysis buffer (Beyotime, Shanghai, China) using a PMSF (Amresco, Solon, Ohio, USA) for 30 minutes at 4 ° C, then centrifuged at 12,000 × g for 15 minutes. 4 ° C. Supernatants were collected and total protein concentration was measured using a BCA Protein Assay Kit (Thermo Scientific, Waltham, MA, USA). An equimolar amount of protein was loaded into each well and separated by 12% SDS-PAGE. The proteins were then transferred to a 0.45-μm PVDF membrane (Amersham Hybond, GE Healthcare, München, Germany) which was blocked in 2% bovine serum albumin (Amresco, Solon, Ohio, USA) and then incubated overnight at 4°. C has the following primary antibodies:: rabbit anti-IL-1RA, rabbit anti-IL-1α (1:1000), mouse anti-β-actin (1:2,000; Cell Signaling Technology, Danvers, MA, USA), rabbit anti-CD31(1:1000, AF6191), rabbit anti-PI3K (1:1000, ab140307), rabbit anti-PI3K alpha(1:1000, ab125633), rabbit anti-NF-κB (1:1000, ab7449), rabbit anti-NF-κB alpha(1:1000, ab194729), and VEGF-A polyclonal antibody (1:1000, A41552). After washing 3 times in TBST buffer for 10 min, the cells were incubated with secondary antibody for 1 h at room temperature. The imprinted material was prepared by chemiluminescence enhancement technique (Xiamen, Lulong Biotechnology).

### 2.4 Vector construction and lentivirus transduction

The open reading framework of the human IL-1RA gene was amplified by PCR-and cloned into a flat viral expression vector, PCDH-CMV-MCS-EFP-Puro(systems biology science, Mountain View, Calif.). pMDL, pVSVG and pREV conversion of recombinant plasmid or empty vector to packaged plasmid into 293T cells. The upper body was collected after transfection at 48H and used to infect kyse410 and Eca109 cells cultured in 6 cm plates. Resistant cloning of puro was expanded into cell lines, such as IL-1RA over-expression cells(KYSE410-IL-1RA or Eca109-IL-1RA), or KYSE410-pCDH or Eca109-pCDH). The protein expression level of IL-1RA was evaluated by Western blotting analysis and RT-PCR method.

### 2.5 Cell proliferation and colony formation assays

Cell proliferation was detected by CCK-8 assay. In logarithmic growth phase, cells were seeded in 96-well plates with a density of 4 × 10^3^ / well. The next day, a cell count kit (CCK-8; Donjido, Kumamoto, Japan) was used according to manufacturer’s instructions. The optical density (OD) was measured by a microplate reader(BioTek, VT, USA) for 6 days a day.

For colony formation assays, 500 cells were inoculated in 6-well plate. 14 days later, 30 min,1% crystal violet staining with 3% methanol was used for counting under inverted microscope for 10 min,. All experiments are in triplicate.

### 2.6 Migration and invasion assays

The system is used to detect cell migration according to the manufacturer’s protocol. For each well, 5×10^4^ cells are sown in the upper chamber of the outflow plate in serum-free RPMI-DMEM media, while 10 % FBS media are added to the lower chamber. The cells left in the upper cavity are scraped off, then fixed in methanol, and stained with 0.1 % of the crystalline purple solution after 48H hatching. Five random fields of view were selected to calculate the cells migrated to the lower side. In the intrusion detection, 10^5^ cells were coated with matrix gel in the upper cavity. The migration test is then carried out.

The transfected cells grew to 100% confluence in 6-well plate. The cell layer was cut at the end of 20 μL to form the wound gap, washed with phosphate buffer solution for three times, and photographed at different time points. Cells were knocked down for 48 hours and counted with a ruler.

### 2.7 in vivo tumor growth study

A total number of 1 ×10^6^ KYSE410 and Eca109 cells which stably overexpressing IL-1RA or control RNA were injected into flanks subcutaneously of BALB/C nude mice (4-week-old male, n=6 per group). The tumor volumes were measured on days 7, 14, 21, 28, 35, and 42 after the implantation procedure. The volume of Tumor was calculated using the equation V (mm3) = length × width. The tumors were collected for detecting weight. All animal studies were reviewed and approved by the Institutional Animal Care and Use Committee of Fujian Medical University.

### 2.8 Elisa for VEGF

Elisas were performed with a commercially available ELISA kit (ab222510, Abcam). The supernatants of cultured KYSE410 and Eca109 cells stably overexpressing IL-1RA or control RNA were collected. Then, VEGF concentrations were detected according to the manufacturer’s instructions.

### 2.9 HUVEC Tube formation assay

Growth factor-reduced Matrigel Basement Membrane Matrix (BD Biosciences, Franklin Lakes, NJ, USA) was slowly thawed on ice, and to each well of a 96-well plate was added 50 ul for polymerization. A number of 1 × 10^4^ HUVECs was plated on top of the Matrigel matrix and treated with the supernatants from KYSE410 and Eca109 cells stably overexpressing IL-1RA or control RNA cultured VEGF for 24 h. The capillary network was analyzed by calculating the length of tubes in 10 random microscopic fields using computer-assisted microscopy.

### 2.10 Statistical analysis

Data were expressed as mean ± standard deviation (SD) of at least three independent experiments performed in triplicate. Statistical analyses was performed in SPSS 23.0 statistical software (SPSS Inc., Chicago, IL, USA). Differences between the two groups were assessed by unpaired Student’s t test, Data with P < 0.05 were considered significant.

## 3. Result

### 3.1 IL-1 overexpression inhibits proliferation and migration of ESCCcells in vitro

We constructed IL-1 overexpressing cell lines in esophageal squamous carcinoma cells KYSE410 and Eca109 by lentiviral transfection. The expression of IL-1RA was verified by WB and q-PCR (Fig 1. A and Fig 1. C). Then, the effect of IL-1RA working on the proliferation of esophageal squamous carcinoma cells was examined by cell plate cloning assay and CCK-8 cell proliferation assay in vitro. The results showed that overexpression of IL-1RA can significantly reduce the proliferation of esophageal squamous carcinoma cells, (p < 0.001; p < 0.05)(Fig 2. A and Fig 2. B). Secondly, we examined the effect of IL-1RA on the migration of esophageal squamous carcinoma cells by cell Transwell chamber assay. The results showed that overexpression of IL-1RA can significantly reduce the migration of ESCC. P<0.0001(Fig 2. C).

**Fig 1.**
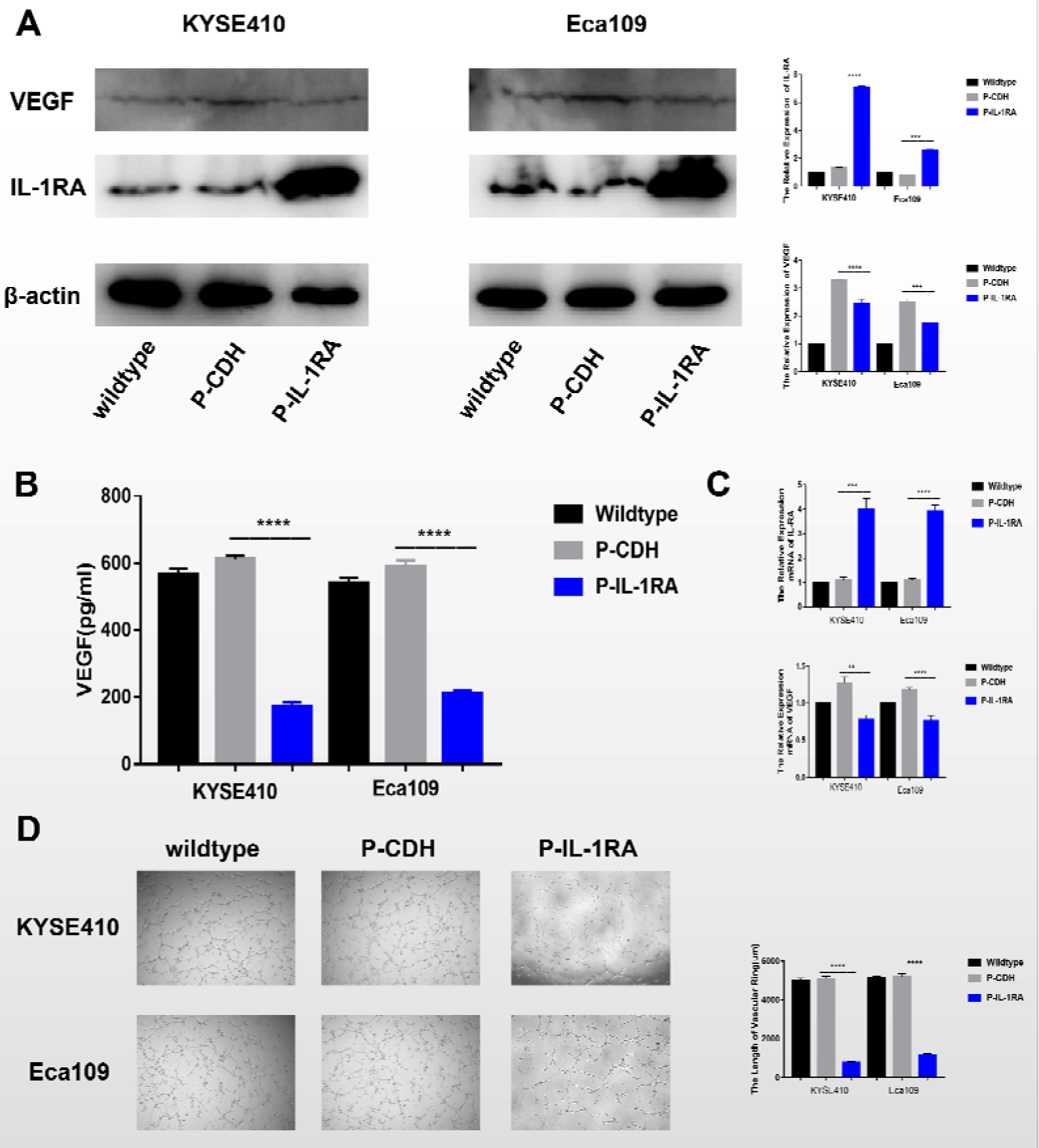
Overexpression of IL-1RA inhibits the expression and secretion of VEGF and inhibits tumor angiogenesis. (A) Western blot analysis confirming expression of IL-1RA in ESCC, and the expression of VEGF. (B) Elesa was used to detect the secretion of VEGF in cell supernatant. (C) RT-PCR was used to confirming expression of IL-1RA in ESCC, and the expression of VEGF.(C) HUVEC Tube formation assay.

**Fig 2.**
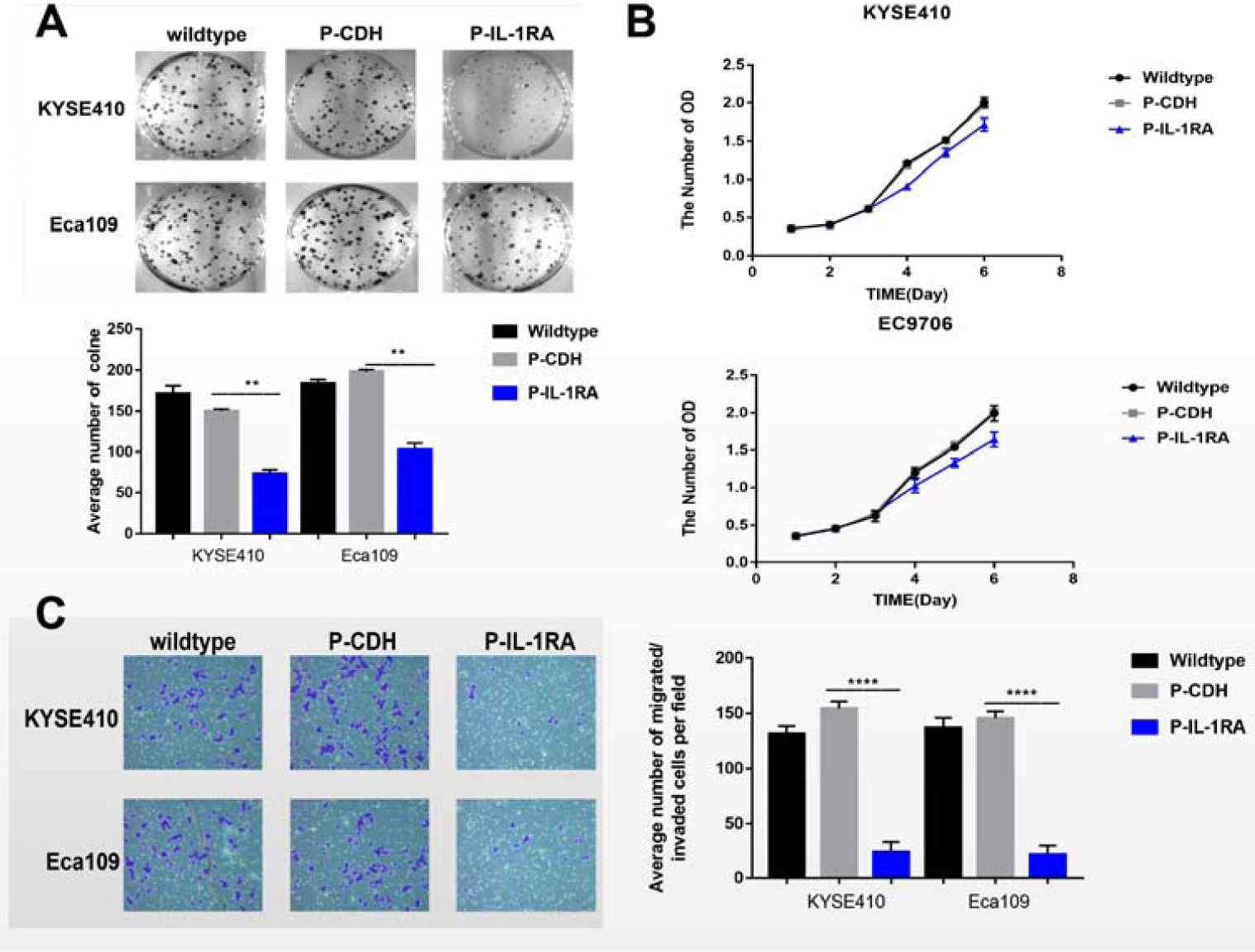
IL-1RA regulated GC cell proliferation and migration. (A) Colony formation assay. (B)CCK-8 assay.(C)Relative migration of the GC cells through an uncoated filter toward serum-containing medium in a Boyden chamber assay.

### 3.2 Overexpression of IL-1RA inhibits the expression and secretion of VEGF in vitro

We expressed the expression of VEGF in ESCC stably overexpressing IL-1RA and its control strain by WB and ELESA. The results showed that overexpresion IL-1RA were expressed in esophageal squamous carcinoma cells KYSE410 P-IL-1RA and Eca109. the expression of VEGF was lower in the ESCC over-expression IL-1RA than the control group (Fig 1. A and Fig 1. c), while the secretion of VEGF in the supernatant of elesa assay showed that the secretion of VEGF in overexpressing IL-1RA cells was significantly reduced (Fig 1. B). The HUVEC Tube formation assay of 6 groups of cell supernatants also showed that the tube formation ability of IL-1RA over-expression group was significantly down-regulated (Fig 1. D), which also suggested that IL-1RA can reduce the expression and secretion of VEGF.

### 3.3 Overexpression of IL-1RA inhibits tumor proliferation and angiogenesis in vivo

We took the ESCC KYSE410 P-IL-1RA, KYSE410 P-CDH, Eca109 P-IL-1RA and Eca109 P-IL-1RA for subcutaneous tumor formation in nude mice. The results showed that The KYSE410 P-IL-1RA and Eca109 P-IL-1RA cells were significantly down-regulated in the nude mice regardless of tumor formation rate or tumor formation volume with time (Fig 3. A, 3. B and 3. C). The WB results from subcutaneous tumor-forming tissue showed that the expression of angiogenic protein CD31 was also down-regulated in the tumor-bearing tumors of the overexpressed IL-1RA group (Fig. 3. D).

**Fig 3.**
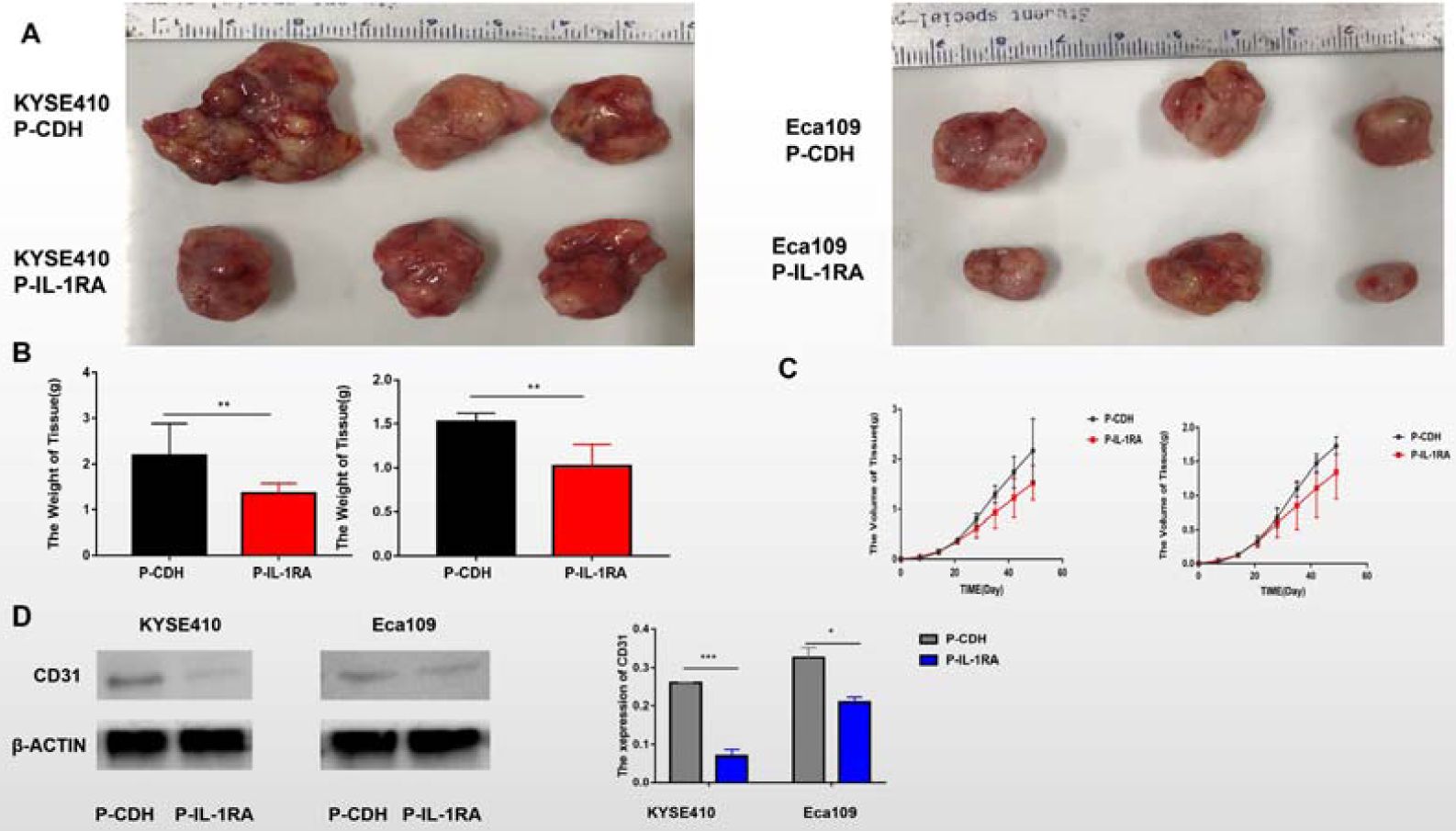
Overexpression of IL-1RA significantly inhibits EC tumor growth and angiogenesis in nude mice in vivo.(A) Tumor growth in nude mice 25 days after injection ESCC. (B)The weigh of tumor tissue from nude mice after injection ESCC.(C) Time-dependent growth of xenograft tumor tissues in nude mice. (D)Altered expression of CD31 in dissected tumor tissues.

**Fig 4.**
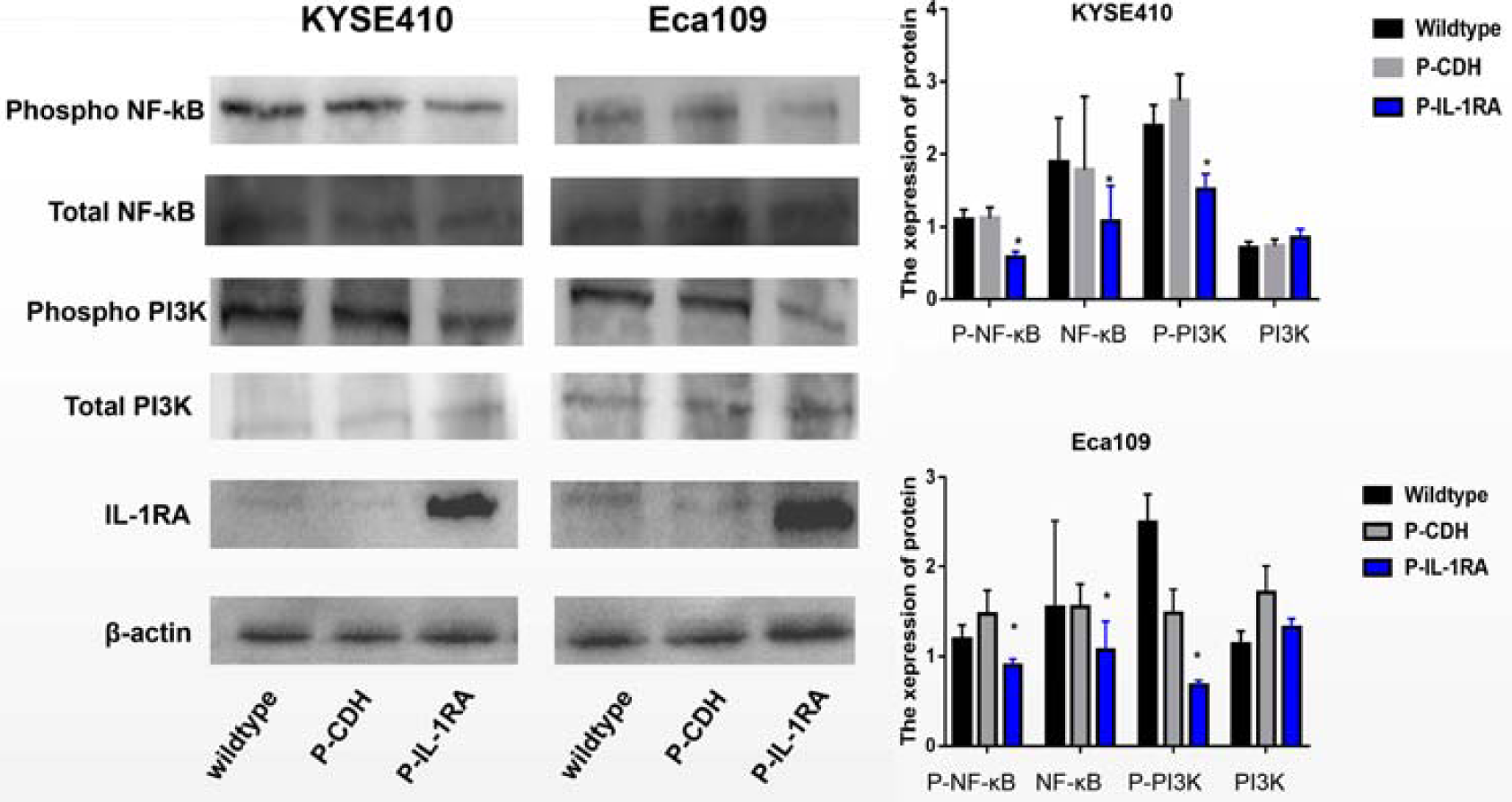
Regulation of VEGF by IL-1RA was dependent on PI3K/NF-κB pathway. Western blot analysis of PI3K, p-PIK, NF-κB, p-NF-κB expression in indicated esophageal cancer cell lines.

### 3.4 VEGF expression may be regulated by PI3K/NF-κB signaling pathway In ESCC

We examined the expression of PI3K and its phosphorylated forms, NF-κB and its phosphorylated forms in esophageal squamous carcinoma cells with wild-type, overexpressing IL-1RA and its control group by WB. The results showed that overexpression of IL-1RA was observed. The total amount of PI3K did not change significantly, but its phosphorylated form was significantly reduced, suggesting that PI3K activity was inhibited, and NF-κB downstream of its signaling pathway was down-regulated regardless of total expression or phosphorylation activation pattern (Fig 3). These results suggest that IL-1RA may regulate tumor angiogenesis by regulating PI3K/NF-κB regulation of VEGF expression and inhibit tumor progression.

## 4. Discussion

The results of the studies reported here provide the evidence that the expression of IL-1RA can effect the biological behavior of EC. In the cells experimental model, stabe over-expression of IL-1RA in the ESCC decreased proliferation and migration. The over-expression of IL-1RA in the ESCC can Reduce the expression and secretion of VEGF through inhibition of PI3K/NF-κB signaling pathway. Combined with our previous experimental results, IL-1RA is lowly expressed in esophageal cancer tissues and is associated with high TNM staging and poor survival prognosis. We can implicate that IL-1RA functions as a tumor suppressor and that its loss can promote the expression and secretion of VEGF and tumor progression.

Angiogenesis is a physiological process in which neovascularization is gradually formed from a portion of a blood vessel pre-existing. In 1971, Dr. Judah Folkman verificated that tumor growth is dependent on the angiogenesis. He found that the diffuse substances secreted by the tumor can stimulate the proliferation of endothelial cells, and then form new blood vessels, and the tumor must form new blood vessels to provide nutrients after growing for more than a few millimeters[18]. It has been commal accepted that VEGF-A is the earliest discovered and most important pro-angiogenic factor[19]. It has been demonstrated that VEGF overexpression potentiates the migratory and invasive ability of ESCC cells[20], which coincides with observations in other malignancies[21][22]. Our experimental results show that the overexpression of IL-1RA in ESCC can reduce the expression and secretion of VEGF, and inhibiting angiogenesis in tumors. In animal experiments, CD31, a marker of vascular formation in the subcutaneous tumor-forming, protein extracted from nude mice, was also significantly reduced in the overexpression of IL-1RA group, suggesting that IL-1RA can inhibit tumor angiogenesis and inhibit tumor progression in vivo.

The formation of the cancer is a multi-step process that include initiations, promotions and progressions. Many pro-inflammatory mediators, involving IL-1, TNF, IL-6, stimulate intracellular kinases Phosphorylating, induce transcription factors such as NF-κB and AP-1, and early response of genes. Which is same to the mode of the classical chemical tumor promoters, TPA and Okadaic acid, supporting cell growth and proliferation via activating cell signaling[23]. Early studies have confirmed a positive correlation between IL-1β secretion in gastric cancer and NF-κB activation in tumors[24]. Another study also reported that infection with H. pylori and IL-1β deficient mice does not activate gastric mucosal inflammation and NF-κB in epithelial cells[25]. IL-1RA inhibits IL-1’s tumor-promoting function by blocking IL-1 binding to its receptor[26]. Our results show that overexpression of IL-1RA can reduce the phosphorylation level of PI3k in cells and further reduce the expression of NF-κB and it’s phosphorylation, suggesting that IL-1RA may play a role through the PI3K/NF-κB signaling pathway.

Study has reported that IL-1 can express the retinoid x receptor alpha expression of downstream proteins induced by NF-κB[27]. And NF-κB can promotes cardiomyocyte growth via binding to the two NF-κB recognition which site in the promoter of VEGF [28]. Therefore, we hypothesized that IL-1RA may inhibit the binding of IL-1 to its receptor, thereby inhibiting the PI3K/NF-κB signaling pathway, reducing the expression and secretion of VEGF, inhibiting the formation of tumor blood vessels and Inhibiting the development of ESCC.therefore, IL-1RA might serve as a potential molecular target for EC treatments.

## Acronyms

IL-1RA: Interleukin 1 receptor antagonist
IL-1β: Interleukin-1β
IL-1RN: Interleukin 1 receptor antagonist gene
VEGF: Vascular endothelial growth factor
ESCC: Esophageal cancer cells
PI3K: Phosphatidylinositol 3-kinase
NF-κB: Nuclear factor-kappa B

